# Inside-Out: Modeling the link between Zika virus viral dynamics within hosts and transmission to vectors across host species and virus strains

**DOI:** 10.1101/2025.04.23.650178

**Authors:** Hélène Cecilia, Benjamin M. Althouse, Sasha R. Azar, Shannan L. Rossi, Nikos Vasilakis, Kathryn A. Hanley

## Abstract

Epidemiological models of mosquito-borne virus transmission often lack accurate estimates of host-to-vector transmission probability. Here, we estimated this probability for two strains of Zika virus (ZIKV)—one sylvatic and one human-endemic—from two monkey species to *Aedes albopictus* mosquitoes using experimental infection data. Viral dynamics did not differ between monkey species, although one (cynomolgus macaque) is a native ZIKV host and the other (squirrel monkey) a novel host, but differed between strains, with viremia for the human-endemic strain peaking later and lower than the sylvatic strain. Only the sylvatic strain was transmitted to mosquitoes. In mosquitoes, anatomical barriers influence viral progression to salivary glands, complicating host infectiousness estimation. We quantified the probability of viral dissemination to the legs in *Ae. albopictus*, which increased with host viral load and was higher after feeding on squirrel monkeys than on cynomolgus macaques. We also found a positive relationship between virus titer in mosquito legs and virus detection in saliva after a 14-day extrinsic incubation period. Combining these factors, we found that squirrel monkeys were on average 1.5 times more infectious to *Ae. albopictus* than cynomolgus macaques. These estimates will help assess ZIKV’s potential to establish an enzootic, sylvatic cycle in the Americas.

## Introduction

Understanding the factors that influence pathogen transmission is essential for predicting and controlling the spread of infectious diseases. In particular, estimating infectiousness—the probability that an infected host transmits a pathogen—is crucial for developing accurate epidemiological models [1]. Modeling at the within-host scale has been employed to derive infectiousness estimates of SARS-CoV-2 and influenza in humans [2], foot-and-mouth disease virus in cattle [3], and influenza virus in pigs [3], among others. For vector-borne pathogens, the infectiousness of one host to the next also depends on the vector that bridges the two hosts, as different vector taxa can vary in their competence. Infectiousness estimates exist for dengue virus (DENV) from humans to *Aedes aegypti* mosquitoes [4], Rift Valley fever virus from cattle, sheep, and goats to *Aedes* spp. and *Culex* spp. mosquitoes [5], Usutu virus from birds to *Culex quinquefasciatus* mosquitoes [6], and Zika virus (ZIKV) from humans to *Aedes albopictus* mosquitoes [7].

For mosquito-borne viruses, it is commonly assumed that within-host viral load is positively related to the probability of host-to-vector transmission [8], although some counter examples have been observed (e.g. DENV in [9]). In the vector, a virus must overcome sequential anatomic barriers (i.e, the midgut infection barrier, the midgut escape barrier, and the salivary gland infection and escape barrier) before becoming infectious [10,11]. Experimental constraints can make it difficult to gather enough information to characterize host infectiousness with regard to capacity to generate saliva-positive vectors over the complete course of host infection. Hence, existing estimates of host infectiousness are often limited to vector infection or disseminated infection [4,5,7]. Besides, transmission experiments sometimes deliver virus to hosts via needle rather than the bite of an infected vector [5,6], or use artificial blood meals to infect vectors [7], which makes their infectiousness estimates less reflective of the natural transmission arc.

The current study focuses on Zika virus, which originated in a sylvatic cycle involving non-human primate (NHP) hosts and arboreal *Aedes* mosquitoes in Africa [12,13]. It eventually spilled over into humans, establishing human-endemic transmission in the paleotropics mediated by *Ae. aegypti* and *Ae. albopictus* [14]. In addition to its role in epidemics, *Ae. albopictus* may also contribute to both spillover and spillback, as a bridge vector [15,16]. ZIKV reached the Americas early in the 2010s, resulting in widespread human infections [17]. It is not yet known whether the virus will establish new sylvatic cycles within American NHP populations [13,18,19]. Should it do so, control of ZIKV in the Americas will become more complicated and eradication will likely become impossible. In this context, it is crucial to explore the potential for novel monkey hosts to sustain ZIKV transmission. To this end, we aimed to compare the infectiousness of native and novel monkey host species, resulting from the coupling between their within-host viral dynamics and the associated transmission to *Ae. albopictus*, following experimental infection with ZIKV strains derived from sylvatic [9] and human-endemic cycles (this paper).

Here, we used these data for phenomenological modeling at the within-host scale to characterize viral load dynamics of ZIKV, focusing on the impact of monkey species, viral strain, initial viral dose received, and inter-individual heterogeneity. We then described host-to-vector transmission occurring during those experimental infections, using dose-response relationships, for disseminated infection and saliva infection of *Ae. albopictus*. Finally, we estimated the infectiousness profile of native and novel monkey hosts over the course of their infection, which can readily be used by models of ZIKV transmission dynamics at the population scale.

## Material and Methods

### Ethics statement

Our study complies with all relevant ethical guidelines and all community standards for containment of infected arthropod vectors; all procedures conducted on non-human primates were approved via UTMB Institutional Animal Care and Use Committee (IACUC) protocol 1912100, approved on December 1, 2019.

### Overview of the experiment

Here we leverage data from previous experimental infections of 3 (1M, 2F) adult cynomolgus macaques (*Macaca fascicularis*) and 10 (5M, 5F) adult squirrel monkeys (*Saimiri boliviensis*) with sylvatic Zika virus strain DakAr 41525 (GenBank accession number EF105379.1) delivered by the bites of batches of 15 *Ae. albopictus* mosquitoes, described in Hanley et al. 2024 [9] (Figure S1). Infecting mosquitoes had been inoculated intrathoracically with the virus [9]. We also leverage data from another arm of the study (Figure S1), not previously reported, in which 4 adult cynomolgus macaques (2 males, weighing 4.7 and 5.2 kg at the start of the experiment, and 2 females, 2.8 and 2.9 kg at the start of the experiment) were infected using identical methods with human-endemic ZIKV strain PRVABC-59 (passaged six times in Vero cells, GenBank accession number KU501215). The number of mosquitoes salivating virus and the total dose of virus delivered was estimated by forced salivation conducted 2 days after infected mosquitoes fed on monkeys; the values for ZIKV DakAr 41525 and PRVABC-59 can be found in Supplemental Table S1.

Quantification of viremia from serum as well as feeding of uninfected *Ae. albopictus* mosquitoes on each animal was done regularly between infection and 28 days post-infection [9] (Figure S1). Viral load was measured by serial dilution of sera which were used to infect Vero cells (CCL-81) followed by immunostaining to yield measurements in plaque forming units (PFU, [9]). Engorged mosquitoes from all feeding days were tested for presence of virus in their bodies and legs, 14 days after feeding (unique extrinsic incubation period (EIP) tested), as described in [9]. Mosquito saliva was collected and tested only for those mosquitoes that fed on days 3 and 4 (sylvatic strain) or 3, 4, and 5 (human-endemic strain).

The viremia of monkeys over time can be found in Supplemental Table S2 for the ZIKV sylvatic strain and Supplemental Table S3 for the ZIKV human-endemic strain. Of the four cynomolgus macaques infected with human-endemic ZIKV (FR1565), one is excluded from the analyses presented in this paper as it never became detectably viremic nor transmitted to mosquitoes. Data on transmission to mosquitoes can be found in Supplemental Table S4, for ZIKV sylvatic strain only. Indeed, we only detected one disseminated infection in mosquitoes for the human-endemic strain of ZIKV, and no positive saliva, so we did not analyze this dyad further for host infectiousness.

### Within-host viral dynamics

To describe within-host viral dynamics, we used a phenomenological model adapted to capture the pattern of acute, short-lived infections, which relied on the simple equation [3,20] :

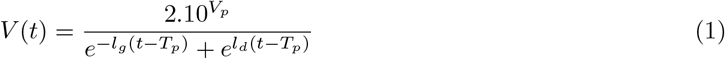

where viral load *V* at time *t* is expressed as a function of peak viral load *V*_*p*_ on a log_10_ scale, time of peak *T*_*p*_, and rates of the exponential growth (*l*_*g*_) and decay (*l*_*d*_) phases. We fitted this equation to viral load data from all monkeys simultaneously, using a non-linear mixed effect modeling approach. We allowed all four parameters to vary with monkey species, viral strain, and dose delivered to the monkeys (fixed effects on three covariates) as well as between individuals (random effects). We used the stochastic approximation expectation maximization (SAEM) algorithm, implemented in the R package saemix, version 3.3 [21]. Parameters were assumed to follow a log-normal distribution, meaning that parameters *θ*_*i*_ of individual *i* can be expressed as *θ*.*e*^*f*^*i*, with *θ* representing the population-level estimate, and *f*_*i*_ = *c*_*i*_.*β* + *η*_*i*_, with *c*_*i*_ representing the subject-specific covariates, *β* the fixed effects, and *η*_*i*_ the random effects, which follow a multinormal distribution *η* ~ *N* (0, *ω*). In addition, we used a constant residual error model. Data below the limit of detection of the assay was set arbitrarily as half the limit of detection if transmission to mosquitoes was detected, 0 otherwise. Further details about the fitting procedure and the computation of uncertainty can be found in Supplemental text S1.

### Vector infection

As mosquito saliva was only tested on a subset of days, we could not estimate host infectiousness over their whole infection period in a single step. Instead, we first analyzed the drivers influencing ZIKV disseminated infection in mosquitoes on all days post infection (dpi), and we then explored the relationship between disseminated infection and ZIKV presence in mosquito saliva, to ultimately infer the whole dynamics of host infectiousness.

### Disseminated infection

We investigated the combined effects of host viral load, host species, and dpi, on the probability of disseminated infection in vectors (*P*_*leg*_). This probability was defined as the ratio of virus-positive legs over the total number of mosquitoes tested. Sometimes, the ratio of virus-positive legs over the number of positive mosquito bodies is used to measure dissemination [22], but in our case an important proportion of mosquito legs were positive when the corresponding bodies were negative [9], which made this ratio uninformative. We performed a model selection procedure similar to what was conducted by Lambrechts et al. [23], to allow non-linear effects (Supplemental text S2.1). A similar approach was used to model the value of the virus titer measured in mosquito legs (*V*_*leg*_).

### Probability of ZIKV presence in saliva

We then explored the relationship between *V*_*leg*_ and the probability of ZIKV being detected in that mosquito’s saliva (*P*_*saliva*_), both measured after the same EIP. To do so, we selected among three functional forms with a sigmoidal shape, and between the use of a binomial or a betabinomial likelihood, the latter accounting for overdispersion in the data ([9], Supplemental text S2.2). The selection was done based on the corrected Akaike Information Criterion (AICc, the lowest AICc was selected).

### Host infectiousness over time

We expressed host infectiousness over time as variations of the transmission rate which, in vector competence studies, is defined as the proportion of saliva-positive mosquitoes among all mosquitoes tested. To do so, we combined the different steps presented above, with the associated uncertainties, as presented in Figure S2.

## Results

### Within-host viral dynamics

Dose delivered to the monkeys had no significant effect on the parameters of Eq. 1 (Table S1). Below, we present the results of a model including only monkey species and viral strain as covariates.

Viral strain had a significant effect on *V*_*p*_ and *T*_*p*_ (Table 1). Indeed, in macaques, the human-endemic strain of ZIKV induced significantly lower and later peak titers than the sylvatic strain (Figure 1 C,E). Monkey species had no significant effect, as macaques and squirrel monkeys infected with the sylvatic strain showed similar viral dynamics (Figure 1 A,C).

**Table 1:**
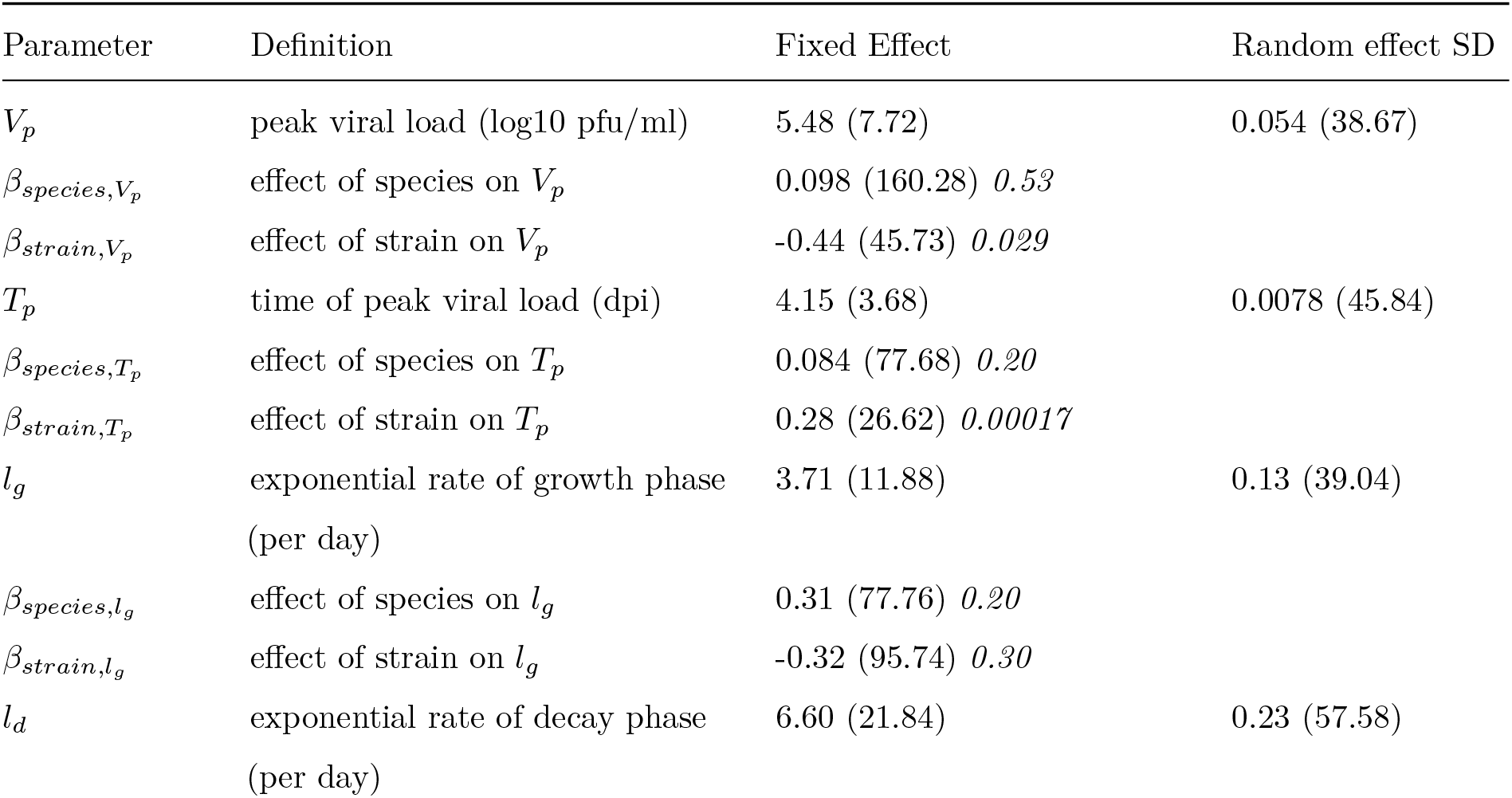
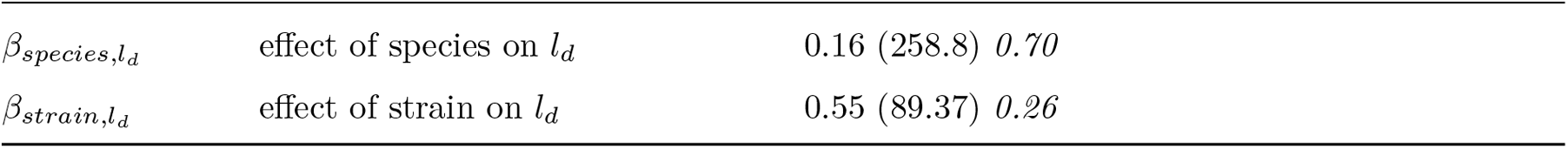
Results of the non-linear mixed effect model of within-host viral dynamics, testing the effect of monkey species and viral strain. Parameter estimates (relative standard error, %) *p value* (for covariates species and strain only). Squirrel monkeys infected with the sylvatic strain of ZIKV are taken as the reference. SD: standard deviation, dpi: days post infection.

**Figure 1:**
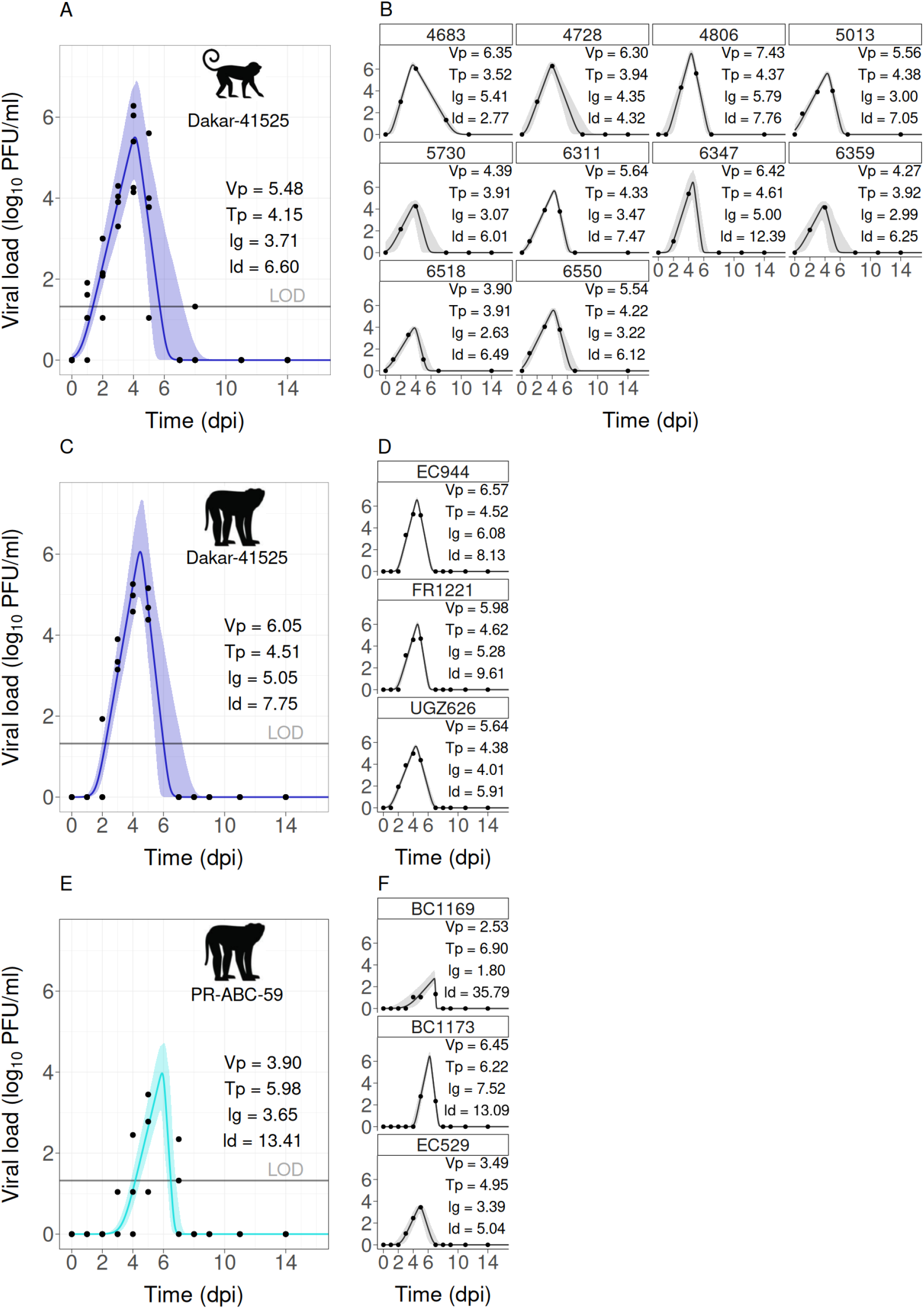
Data and model fit for within-host viral dynamics. A-B: squirrel monkeys infected with sylvatic ZIKV. C-D: cynomolgus macaques infected with sylvatic ZIKV. E-F: cynomolgus macaques infected with human-endemic ZIKV. Left column (A,C,E) shows the group-level model fit (line) and associated parameters. The shaded band is the confidence interval of the fixed effects. Right column (B,D,F) shows the individual fits, uncertainty, and associated parameters. Points are raw data. As no transmission to mosquitoes was detected with the hu7man-endemic strain in cynomolgus macaques, we do not analyze this dyad further for host infectiousness. LOD = limit of detection.

There was very little variability among the three macaques infected with the sylvatic strain (Figure 1 D) and more inter-individual variability among the two other groups (Figure 1 B,F). *l*_*d*_ and *T*_*p*_ were the most and least variable parameters, respectively (Table S2).

### Vector infection

#### Disseminated infection

The model selected to describe *P*_*leg*_ included a non-linear, non-decreasing effect of viremia, a non-linear, non-monotonic effect of day post infection, and an effect of monkey species (Figure 2, Supplemental text S2.1.1). Viral load had a significant effect (p = 9.5e-12, Figure 2 A,B). Monkey species had a significant effect (p = 2.5e-5), with transmission from squirrel monkeys being more efficient than from cynomolgus macaques (OR = 4.20, 95% CI [2.15;8.18], Figure 2 A,B).

**Figure 2:**
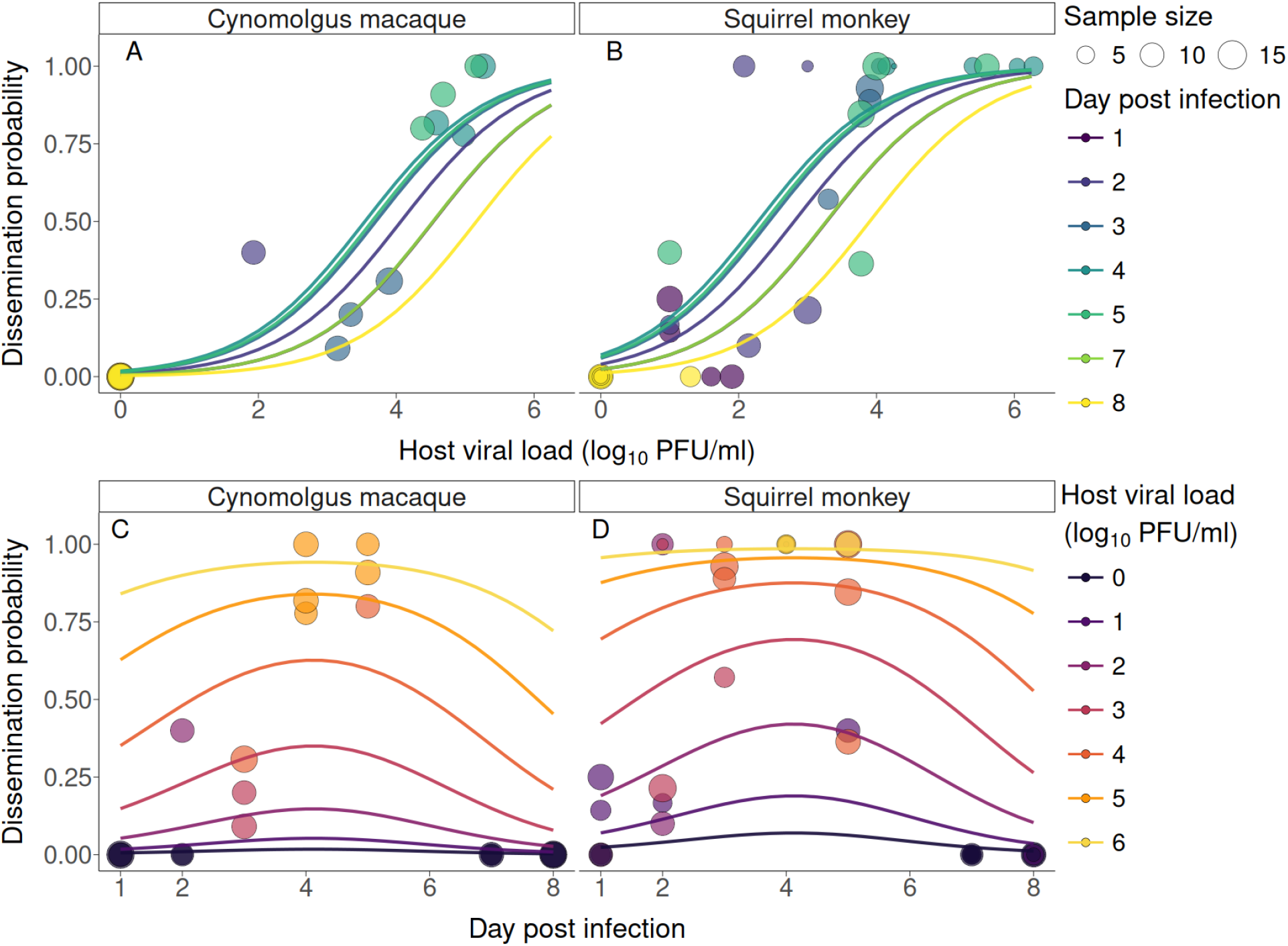
Probability of ZIKV disseminated infection in Ae. albopictus as a function of viral load (A,B), day post infection (C,D), and monkey species (A,C; B,D). Each circle represents a single batch of mosquitoes fed on a given host on a given day, and the size of the circle is proportionate to the number of mosquitoes analyzed. Lines represent the regression fits to the data. Lines and data are color coded according to day post infection (A,B) or host viremia (C,D). In D, note that several points are overlapping for days 4 and 5 and dissemination probability of 1, corresponding to viremia ranging between [4;6] log_10_ PFU/ml.

Although day post infection did not have a significant effect (p = 0.065), an interesting pattern emerged, with *P*_*leg*_ possibly maximized at 4 dpi even when controlling for viremia (Figure 2 C,D). We note that the significance of dpi varied from one model to the next in our selection process, and even among models with a non-linear, bell-shaped effect of dpi, 2 out of 4 models tested estimated the effect of dpi to be significant.

A simple linear model was selected to describe *V*_*leg*_ as a function of host viremia, day post infection, and monkey species (Supplemental text S2.1.2). Viremia was the only variable with a significant effect (p = 7.6e-10). We noticed a strong heterogeneity of leg titers, particularly when host viral load was between 4 and 5 log_10_ PFU/ml, for both monkey species (Figure S3A). This heterogeneity would not be sufficiently captured in the selected model, as viremia only explained 25% of the variance. To remain simple, for the estimation of host infectiousness, we decided to sample *V*_*leg*_ in the whole distribution of leg titers obtained in our experiment. This distribution was bimodal, with frequent sampling around 2.5 and 5.7 log_10_ PFU/ml (Figure S3B).

### Probability of ZIKV presence in saliva

The relationship between *V*_*leg*_ and *P*_*saliva*_ was best described using a logistic equation with binomial likelihood (Eq. S1), with *log*(*β*_0_) = 3.47 [2.75; 4.04] and *β*_1_ = 0.85 [0.56; 1.18] (Figure 3). This relationship corresponds to mosquito infections arising from both monkey species. Since the distribution of leg titers was similar regardless of the transmitting species (Figure S3C) we fitted the relationship for both species combined.

**Figure 3:**
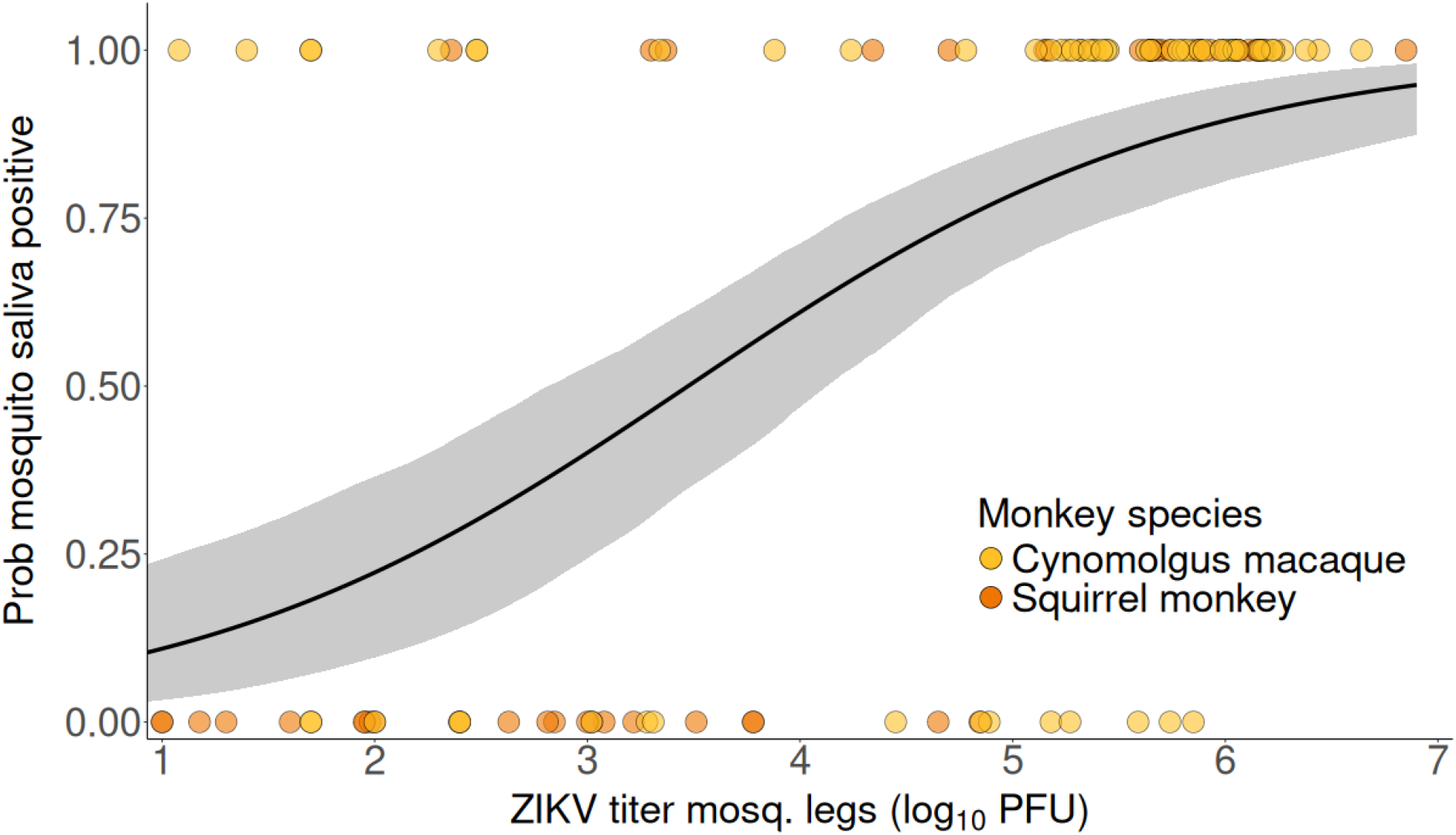
Relationship between ZIKV titer measured in mosquito legs, and probability of mosquito saliva testing positive for ZIKV. Points show raw data, colored by monkey species, each point representing a single mosquito. The line shows the best fit, obtained with Eq. S1 and a binomial likelihood. The shaded band shows the associated 95% confidence interval.

### Host infectiousness over time

The combination of the different steps resulted in the estimation of transmission rates in *Ae. albopictus* over time when feeding on each monkey species infected with sylvatic ZIKV. Squirrel monkeys were overall more infectious than cynomolgus macaques, mainly because their transmission rate increased earlier in infection (Figure 4). This was due to viral loads reaching levels above LOD one day earlier in squirrel monkeys than cynomolgus macaques (Figure S4A), and further exagerated by more efficient dissemination in *Ae. albopictus* after feeding on squirrel monkeys than cynomolgus macaques (Figure S4B).

**Figure 4:**
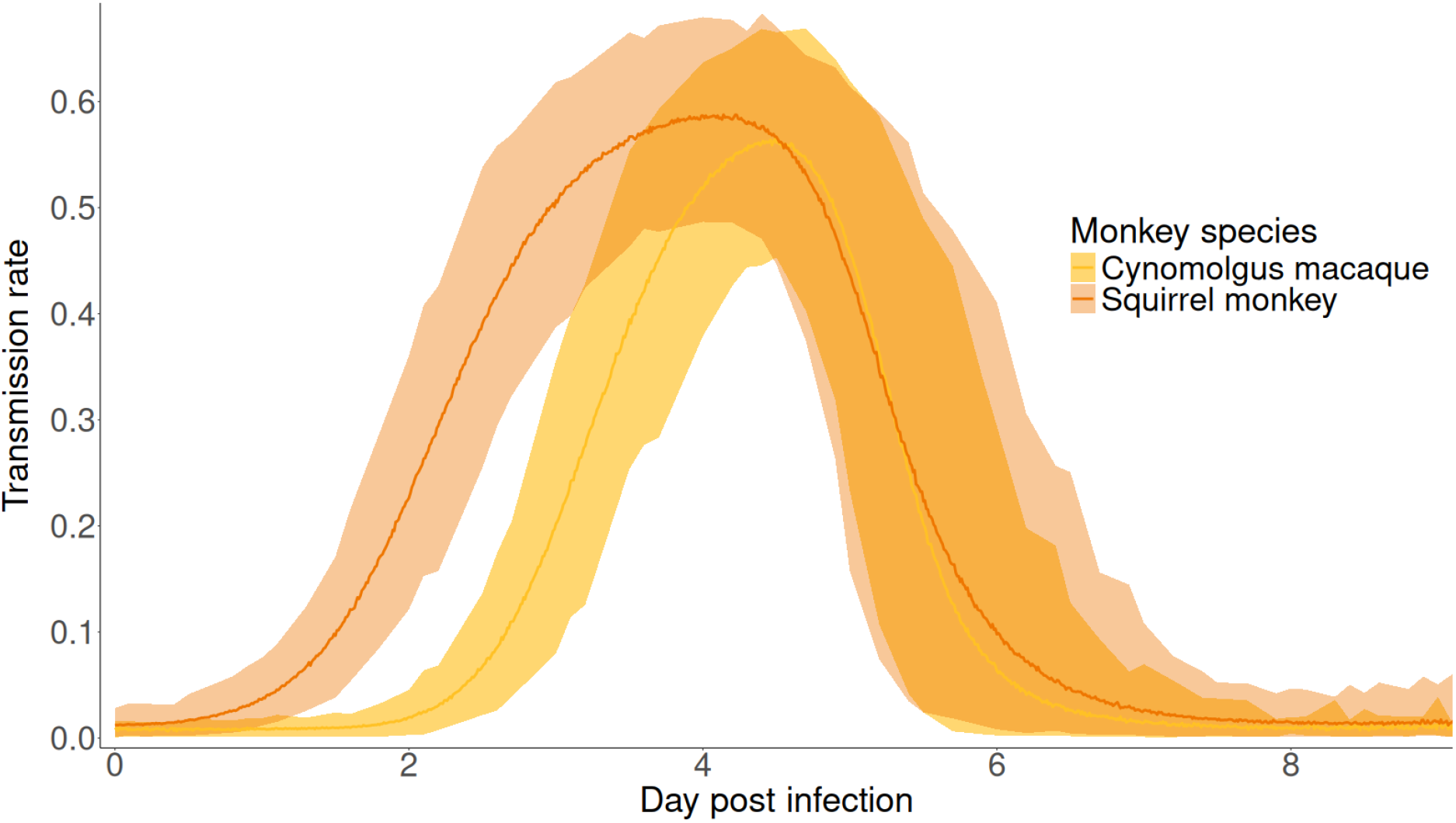
Infectiousness of squirrel monkeys and cynomolgus macaques towards Ae. albopictus over time. Lines show the mean transmission rate (probability of mosquito saliva testing positive for ZIKV over the total number of mosquitoes fed) and shaded bands are the associated 95% confidence interval.

Comparing the area under the curve (AUC) of their infectiousness profiles, we estimated that squirrel monkeys were on average 1.53 times more infectious (95% confidence interval [1.42; 1.63]) than cynomolgus macaques. This is assuming both species are subject to similar contact rates with vectors. The maximum transmission rate was similar for both species (mean 0.56 [0.45; 0.66] for cynomolgus macaques, 0.59 [0.49; 0.68] for squirrel monkeys, Figure 4), due to a common sampling distribution for *V*_*leg*_ and *P*_*saliva*_. This maximum was reached at 4 dpi (Figure 4). As a conservative estimate, we also derived the infectiousness profiles assuming viral loads below LOD would not give rise to any infection (Figure S5). The ratio of AUC between species remained similar (1.49 [1.39; 1.59]).

## Discussion

In this study, we leveraged data from an experiment coupling ZIKV infection of monkeys with transmission to *Ae. albopictus* vectors to derive estimates of monkey infectiousness over time. The experiment was replicated across two species of monkey, one (cynomolgus macaque) a native host of the virus and one (squirrel monkey) a novel host with potential to serve as a New World reservoir in the future, as well as two strains of the virus, one each from the sylvatic and human-endemic ecotype [24]. We found that monkey species had no significant effect on within-host viral dynamics, while virus strain did, with the human-endemic strain inducing later and lower viremia peaks than the sylvatic strain. Moreover, the human-endemic strain failed to transmit to *Ae. albopictus* mosquitoes, but the sylvatic strain robustly transmitted. For the sylvatic strain, host viral load and host species both influenced disseminated infection in mosquitoes (*P*_*leg*_), with more efficient transmission at higher viremia, and from squirrel monkeys than cynomolgus macaques. Turning to infectious mosquitoes, we found a positive relationship between *V*_*leg*_ and *P*_*saliva*_ after an EIP of 14 days. Taken together, these results allowed us to conclude that squirrel monkeys were overall 1.5 times more infectious than cynomolgus macaques towards *Ae. albopictus*, when infected with a sylvatic strain of ZIKV, while the trend in human-endemic ZIKV could not be investigated due to a dearth of transmission.

The transmission rates estimated here can readily be used to improve population-level models, such as Althouse et al. [18]. This underscores the utility of the framework we have developed, which exploits all aspects of the experimental design to provide an accurate picture of monkey infectiousness over time. Indeed, had we only focused on within-host viral dynamics, we would have concluded that squirrel monkeys and cynomolgus macaques were likely to infect mosquitoes similarly, which would have been a mistake. Of note, quantifying viral titer in mosquito legs was instrumental for us to link the different components of infectiousness, but this type of measure is not done systematically in vector competence studies. In the future, our framework can easily be applied to similar experiments involving other arbovirus, hosts, or vectors.

The modeling techniques we used were adapted to the quantity of data available, but do not provide mechanistic insight to explain observed differences between viral strains and monkey species. The equation used to describe within-host viral dynamics has the advantage of only having four parameters, which can all be estimated at the individual and population levels. In comparison, even simple, target-cell limited, mechanistic within-host models, with no compartments for the immune response, which have been used in previous studies of within-host ZIKV dynamics [7,25–27], have 4 to 5 parameters and 2 initial conditions. When these are fitted to datasets of viral loads over time, some of these parameters and initial conditions must be fixed, sometimes arbitrarily or with the help of the literature, which can limit the mechanistic insight gained. When the objective is to capture the overall dynamics to accurately estimate host infectiousness over time, as was the case here, phenomenological modeling is sufficient and easier to implement. If one wishes to dive deeper into the processes of host-pathogen interactions, to explore different immune mechanisms or the effect of treatment, the mechanistic approach may be better suited. This endeavor will however require fitting to data other than viral loads alone [20,28]. Regarding mosquito infection, the development of a dynamic mechanistic model was not possible as we only had data for a unique extrinsic incubation period. Acquiring data at different extrinsic incubation periods usually requires sacrificing mosquitoes, which needs to be accounted for in models and also limits the number of individuals tested per time step. To bypass this issue, possible alternatives consist in monitoring viral shedding in mosquito excreta [29], or having live mosquitoes salivate on sugar pads [30].

The human-endemic ZIKV strain used here (PRVABC59) has been used in previous experimental infections of monkeys, albeit comparisons with our study are limited by differences in monkey species used (rhesus macaques *Macaca mulatta* in [25,27,31–35], cynomolgus macaques in [36,37]), inoculation route (subcutaneous in all except Dudley et al. [31] who used both mosquito bite and needle delivery), or type of viremia measured (genome copies in all except Triplett et al. [32]). Dudley et al. [31] used mosquito (*Ae. aegypti*) bites to deliver PRVABC59 to rhesus macaques, and measured viremia via qRT-PCR. Similar to our results, they observed that viremia peaked at day 5 or 6 pi. In Dudley et al. however, viremia was detected earlier (from day 1 pi for 3/4 individuals *vs* never before 3 dpi in our case) and lasted longer (at least until day 8 *vs* never after day 7 in our case). This likely reflects differences in detectability and persistence between viral genome copies and infectious virus. Indeed, in Triplett et al. [32], infectious viremia peaked at 2.5-3 log_10_ PFU/ml in rhesus macaques, similar to our study, and was resolved by day 5. Dudley et al. noted that peak viremia was reached 1-2 days later with mosquito-bite inoculation compared to subcutaneous inoculation. We observe the same delay when comparing our results to studies subcutaneously inoculating doses between 3 and 4 log_10_ PFU ZIKV [25,27,35,37], which is similar to the dose we estimated was delivered by mosquitoes in the current experiment (min-max [2.80-4.47] log_10_ PFU ZIKV). Studies using a sylvatic strain closely related to the one we used (Dakar 41524), all infected pregnant rhesus macaques subcutaneously, and measured viremia through genome copies [35,38,39]. Interestingly in a subset of these, Crooks et al. [35] were able to compare viral dynamics between strain PRVABC59 and Dakar 51424 and observed later and lower peak viremia for PRVABC59, similar to what we found, although these differences were not significant in their case.

Regarding the drivers of host-to-vector transmission, we observed a pattern similar to what was recently reported by Lambrechts et al. [23], who found maximum transmission of DENV from humans to *Ae. aegypti* at day 2 post symptom onset, after controlling for levels of viremia. In our case, maximum transmission was reached at 4 dpi, but this association was not significant in the final selected model. The fact that this pattern emerged in our study, where we measured infectious viremia, compared to the data in Lambrechts et al., who measured genome copies, suggests that infectivity of viral particles *per se* is not responsible for this difference in transmission between early and late infection. The mechanistic basis of this phenomenon remains to be elucidated.

The likelihood of ZIKV establishing a sylvatic transmission cycle in the Neotropics cannot be inferred from our data alone. The human-endemic strain tested here belongs to the clade circulating in the New World, and did not transmit well to NHPs in the present study. However, at least one study has reported circulation of African lineage ZIKV in Brazil [40], and sylvatic DENV has been transported multiple times out of both Africa and Asia to other continents [41–43], demonstrating the potential for sylvatic ZIKV to be introduced to the Americas. Furthermore, the sylvatic strain we tested transmitted efficiently between squirrel monkeys and *Ae. albopictus*, but this is only one host-vector combination out of many possible in the field. The current experimental design already provided evidence that trophic preferences are likely to play a role in transmission dynamics, as engorgement rates were on average higher on cynomolgus macaques than on squirrel monkeys [44]. Finally, the co-occurrence of vectors and hosts must be mapped at high granularity, and mosquito blood meals must be analyzed to identify which vectors competent for ZIKV feed on both humans and NHPs and the frequency at which they do so. Such studies would need to be stratified vertically, as the community composition at ground level can greatly differ from that at canopy level [45,46].

## Acknowledgements

This research was funded by grant 1R01AI145918-02 (to K.A.H., N.V., B.M.A., and S.L.R.) and in part by the Centers for Research in Emerging Infectious Diseases “The Coordinating Research on Emerging Arboviral Threats Encompassing the Neotropics (CREATE-NEO)’ ‘grant 1U01AI151807 from the US National Institutes of Health (to N.V. and K.A.H.). Squirrel monkeys acquired from the UT MD Anderson Cancer Center, Michael E. Keeling Center for Comparative Medicine and Research, were supported by the NIH grant 5P40OD010938-36 prior to sale to UTMB.

## References

1. Martin LB et al. 2019 Extreme Competence: Keystone Hosts of Infections. Trends in Ecology & Evolution 34, 303–314. (doi:10.1016/j.tree.2018.12.009)

2. Goyal A, Reeves DB, Cardozo-Ojeda EF, Schiffer JT, Mayer BT. 2021 Viral load and contact heterogeneity predict SARS-CoV-2 transmission and super-spreading events. eLife 10, e63537. (doi:10.7554/eLife.63537)

3. Gubbins S. 2024 Quantifying the relationship between within-host dynamics and transmission for viral diseases of livestock. Journal of The Royal Society Interface

4. Bosch QA ten et al. 2018 Contributions from the silent majority dominate dengue virus transmission. PLOS Pathogens 14, e1006965. (doi:10.1371/journal.ppat.1006965)

5. Cecilia H, Vriens R, Wichgers Schreur PJ, Wit MM de, Métras R, Ezanno P, Bosch QA ten. 2022 Heterogeneity of Rift Valley fever virus transmission potential across livestock hosts, quantified through a model-based analysis of host viral load and vector infection. PLOS Computational Biology 18, e1010314. (doi:10.1371/journal.pcbi.1010314)

6. Heitzman-Breen N, Liyanage YR, Duggal N, Tuncer N, Ciupe SM. 2024 The effect of model structure and data availability on Usutu virus dynamics at three biological scales. Royal Society Open Science 11. (doi:10.1098/rsos.231146)

7. Lequime S, Dehecq J-S, Matheus S, Laval F de, Almeras L, Briolant S, Fontaine A. 2020 Modeling intra-mosquito dynamics of Zika virus and its dose-dependence confirms the low epidemic potential of Aedes albopictus. PLOS Pathogens 16, e1009068. (doi:10.1371/journal.ppat.1009068)

8. Althouse BM, Hanley KA. 2015 The tortoise or the hare? Impacts of within-host dynamics on transmission success of arthropod-borne viruses. Philosophical Transactions of the Royal Society B: Biological Sciences 370, 20140299. (doi:10.1098/rstb.2014.0299)

9. Hanley KA et al. 2024 Trade-offs shaping transmission of sylvatic dengue and Zika viruses in monkey hosts. Nature Communications 15, 2682. (doi:10.1038/s41467-024-46810-x)

10. Azar SR, Weaver SC. 2019 Vector Competence: What Has Zika Virus Taught Us? Viruses 11, 867. (doi:10.3390/v11090867)

11. Weaver SC, Forrester NL, Liu J, Vasilakis N. 2021 Population bottlenecks and founder effects: implications for mosquito-borne arboviral emergence. Nature Reviews Microbiology 19, 184–195. (doi:10.1038/s41579-020-00482-8)

12. Valentine MJ, Murdock CC, Kelly PJ. 2019 Sylvatic cycles of arboviruses in non-human primates. Parasites & Vectors 12, 463. (doi:10.1186/s13071-019-3732-0)

13. Vasilakis N, Weaver SC. 2017 Flavivirus transmission focusing on Zika. Current Opinion in Virology 22, 30–35. (doi:10.1016/j.coviro.2016.11.007)

14. Mayer SV, Tesh RB, Vasilakis N. 2017 The emergence of arthropod-borne viral diseases: A global prospective on dengue, chikungunya and zika fevers. Acta Tropica 166, 155–163. (doi:10.1016/j.actatropica.2016.11.020)

15. Hanley KA, Monath TP, Weaver SC, Rossi SL, Richman RL, Vasilakis N. 2013 Fever versus fever: The role of host and vector susceptibility and interspecific competition in shaping the current and future distributions of the sylvatic cycles of dengue virus and yellow fever virus. Infection, Genetics and Evolution 19, 292–311. (doi:10.1016/j.meegid.2013.03.008)

16. Pereira-dos-Santos T, Roiz D, Lourenço-de-Oliveira R, Paupy C. 2020 A Systematic Review: Is Aedes albopictus an Efficient Bridge Vector for Zoonotic Arboviruses? Pathogens 9, 266. (doi:10.3390/pathogens9040266)

17. Zhang Q et al. 2017 Spread of Zika virus in the Americas. Proceedings of the National Academy of Sciences 114. (doi:10.1073/pnas.1620161114)

18. Althouse BM, Vasilakis N, Sall AA, Diallo M, Weaver SC, Hanley KA. 2016 Potential for Zika Virus to Establish a Sylvatic Transmission Cycle in the Americas. PLOS Neglected Tropical Diseases 10, e0005055. (doi:10.1371/journal.pntd.0005055)

19. Terzian ACB et al. 2018 Evidence of natural Zika virus infection in neotropical non-human primates in Brazil. Scientific Reports 8, 16034. (doi:10.1038/s41598-018-34423-6)

20. Holder BP, Beauchemin CA. 2011 Exploring the effect of biological delays in kinetic models of influenza within a host or cell culture. BMC Public Health 11, S10. (doi:10.1186/1471-2458-11-S1-S10)

21. Comets E, Lavenu A, Lavielle M. 2017 Parameter Estimation in Nonlinear Mixed Effect Models Using saemix, an R Implementation of the SAEM Algorithm. Journal of Statistical Software 80. (doi:10.18637/jss.v080.i03)

22. Wu VY, Chen B, Christofferson R, Ebel G, Fagre AC, Gallichotte EN, Sweeny AR, Carlson CJ, Ryan SJ. 2022 A minimum data standard for vector competence experiments. Scientific Data 9, 634. (doi:10.1038/s41597-022-01741-4)

23. Lambrechts L et al. 2023 Direct mosquito feedings on dengue-2 virus-infected people reveal dynamics of human infectiousness. PLOS Neglected Tropical Diseases 17, e0011593. (doi:10.1371/journal.pntd.0011593)

24. Musso D, Gubler DJ. 2016 Zika Virus. Clinical Microbiology Reviews 29, 487–524. (doi:10.1128/CMR.00072-15)

25. Osuna CE et al. 2016 Zika viral dynamics and shedding in rhesus and cynomolgus macaques. Nature Medicine 22, 1448–1455. (doi:10.1038/nm.4206)

26. Best K, Guedj J, Madelain V, Lamballerie X de, Lim S-Y, Osuna CE, Whitney JB, Perelson AS. 2017 Zika plasma viral dynamics in nonhuman primates provides insights into early infection and antiviral strategies. Proceedings of the National Academy of Sciences 114, 8847–8852. (doi:10.1073/pnas.1704011114)

27. Best K, Barouch DH, Guedj J, Ribeiro RM, Perelson AS. 2021 Zika virus dynamics: Effects of inoculum dose, the innate immune response and viral interference. PLOS Computational Biology 17, e1008564. (doi:10.1371/journal.pcbi.1008564)

28. Hadjichrysanthou C, Cauët E, Lawrence E, Vegvari C, Wolf F de, Anderson RM. 2016 Understanding the within-host dynamics of influenza A virus: from theory to clinical implications. Journal of The Royal Society Interface 13, 20160289. (doi:10.1098/rsif.2016.0289)

29. Fontaine A, Jiolle D, Moltini-Conclois I, Lequime S, Lambrechts L. 2016 Excretion of dengue virus RNA by Aedes aegypti allows non-destructive monitoring of viral dissemination in individual mosquitoes. Scientific Reports 6, 24885. (doi:10.1038/srep24885)

30. Grubaugh ND et al. 2017 Mosquitoes Transmit Unique West Nile Virus Populations during Each Feeding Episode. Cell Reports 19, 709–718. (doi:10.1016/j.celrep.2017.03.076)

31. Dudley DM et al. 2017 Infection via mosquito bite alters Zika virus tissue tropism and replication kinetics in rhesus macaques. Nature Communications 8, 2096. (doi:10.1038/s41467-017-02222-8)

32. Triplett C, Dufek S, Niemuth N, Kobs D, Cirimotich C, Mack K, Sanford D. 2022 Onset and Progression of Infection Based on Viral Loads in Rhesus Macaques Exposed to Zika Virus. Applied Microbiology 2, 544–553. (doi:10.3390/applmicrobiol2030042)

33. Hirsch AJ et al. 2017 Zika Virus infection of rhesus macaques leads to viral persistence in multiple tissues. PLOS Pathogens 13, e1006219. (doi:10.1371/journal.ppat.1006219)

34. Rayner JO et al. 2018 Comparative Pathogenesis of Asian and African-Lineage Zika Virus in Indian Rhesus Macaque’s and Development of a Non-Human Primate Model Suitable for the Evaluation of New Drugs and Vaccines. Viruses 10, 229. (doi:10.3390/v10050229)

35. Crooks CM et al. 2021 African-Lineage Zika Virus Replication Dynamics and Maternal-Fetal Interface Infection in Pregnant Rhesus Macaques. Journal of Virology 95, e02220–20. (doi:10.1128/JVI.02220-20)

36. Koide F, Goebel S, Snyder B, Walters KB, Gast A, Hagelin K, Kalkeri R, Rayner J. 2016 Development of a Zika Virus Infection Model in Cynomolgus Macaques. Frontiers in Microbiology 7. (doi:10.3389/fmicb.2016.02028)

37. Medina LO et al. 2018 A Recombinant Subunit Based Zika Virus Vaccine Is Efficacious in Non-human Primates. Frontiers in Immunology 9, 2464. (doi:10.3389/fimmu.2018.02464)

38. Rosinski JR et al. 2023 Frequent first-trimester pregnancy loss in rhesus macaques infected with African-lineage Zika virus. PLOS Pathogens 19, e1011282. (doi:10.1371/journal.ppat.1011282)

39. Koenig MR et al. 2023 Vertical transmission of African-lineage Zika virus through the fetal membranes in a rhesus macaque (Macaca mulatta) model. PLOS Pathogens 19, e1011274. (doi:10.1371/journal.ppat.1011274)

40. Kasprzykowski JI, Fukutani KF, Fabio H, Fukutani ER, Costa LC, Andrade BB, Queiroz ATL. 2020 A recursive sub-typing screening surveillance system detects the appearance of the ZIKV African lineage in Brazil: Is there a risk of a new epidemic? International Journal of Infectious Diseases 96, 579–581. (doi:10.1016/j.ijid.2020.05.090)

41. Franco L et al. 2011 First Report of Sylvatic DENV-2-Associated Dengue Hemorrhagic Fever in West Africa. PLoS Neglected Tropical Diseases 5, e1251. (doi:10.1371/journal.pntd.0001251)

42. Pyke AT et al. 2016 Highly divergent dengue virus type 1 genotype sets a new distance record. Scientific Reports 6, 22356. (doi:10.1038/srep22356)

43. Pyke AT, Huang B, Warrilow D, Moore PR, McMahon J, Harrower B. 2017 Complete Genome Sequence of a Highly Divergent Dengue Virus Type 2 Strain, Imported into Australia from Sabah, Malaysia. Genome Announcements 5, e00546–17. (doi:10.1128/genomeA.00546-17)

44. Cecilia H, Althouse BM, Azar SR, Moehn BA, Yun R, Rossi SL, Vasilakis N, Hanley KA. 2024 Aedes albopictus is not an arbovirus aficionado when feeding on cynomolgus macaques or squirrel monkeys. iScience 27, 111198. (doi:10.1016/j.isci.2024.111198)

45. Hendy A et al. 2021 Microclimate and the vertical stratification of potential bridge vectors of mosquito-borne viruses captured by nets and ovitraps in a central Amazonian forest bordering Manaus, Brazil. Scientific Reports 11, 21129. (doi:10.1038/s41598-021-00514-0)

46. Ramdarshan A, Alloing-Séguier T, Merceron G, Marivaux L. 2011 The Primate Community of Cachoeira (Brazilian Amazonia): A Model to Decipher Ecological Partitioning among Extinct Species. PLoS ONE 6, e27392. (doi:10.1371/journal.pone.0027392)

