## Supplementary Information for "Inside-Out: Modeling the link between Zika virus viral dynamics within hosts and transmission to vectors across host species and virus strains"

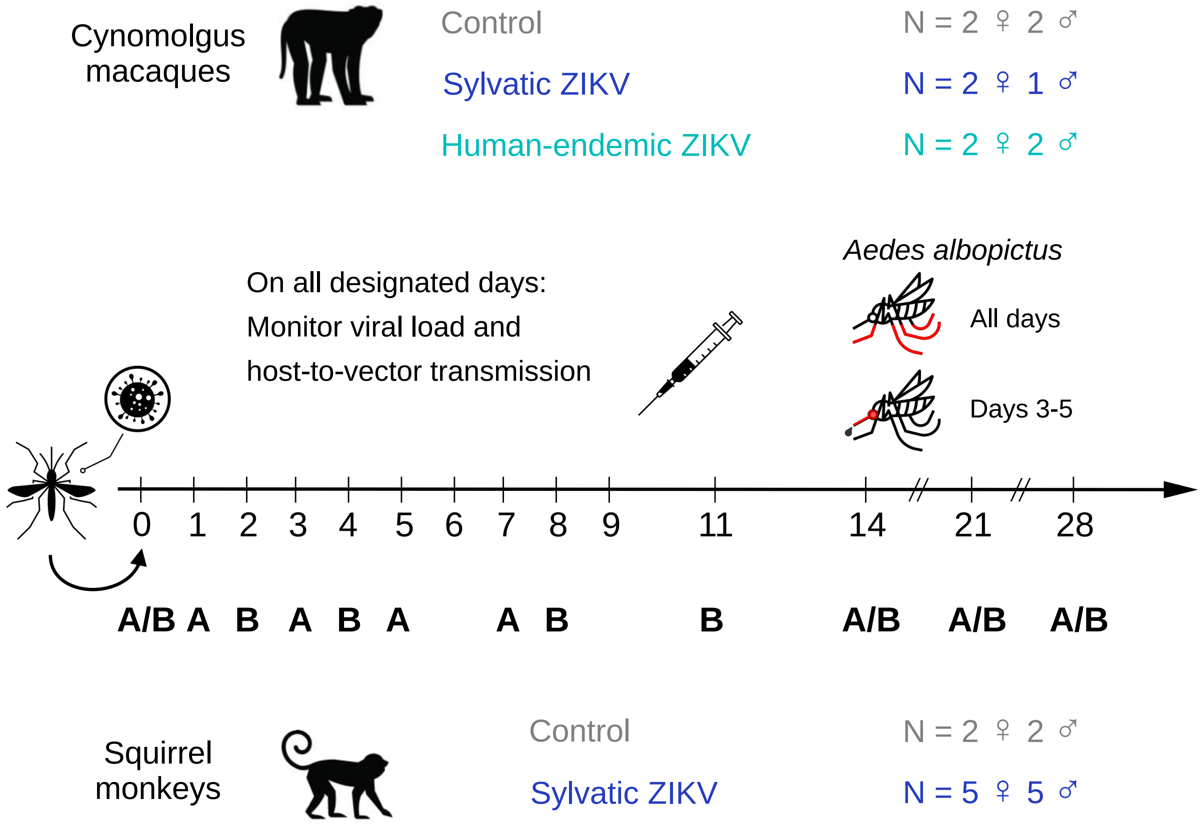


Figure S 1: **Experimental infections of cynomolgus macaques and squirrel monkeys with sylvatic (DakAr 41525) or human-endemic (PRVABC-59) Zika virus.** Monkeys were infected through mosquito bite on day 0. These mosquitoes had been intrathoracically inoculated 10 days before. For mosquitoes used to detect transmission, saliva was tested on days 3 and 4 for the sylvatic strain, and 3, 4, and 5 for the human-endemic strain. Note that 4 cynomolgus macaques were infected with the human-endemic strain, but one never became detectably viremic and is therefore not present in our dataset. A and B indicate cohorts, in the case of squirrel monkeys. The monkey images are licensed from Shutterstock. The mosquito and syringe images were free from Flaticon (see Credits section).

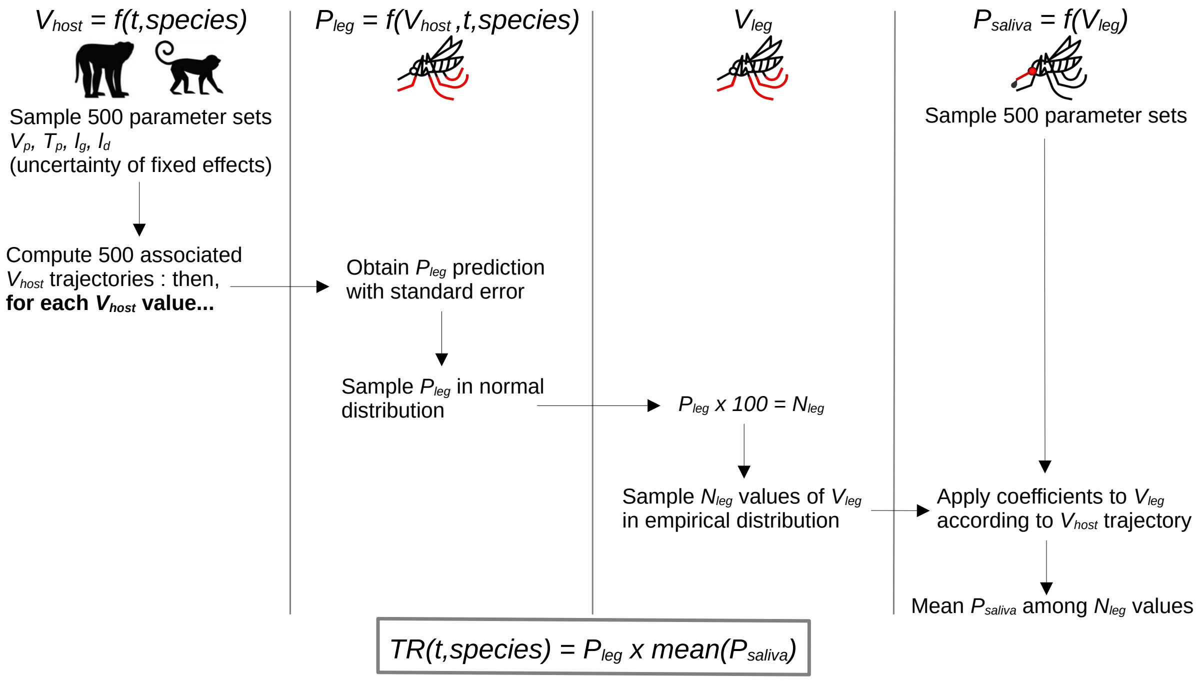


Figure S 2: **Procedure used to compute host infectiousness, defined as transmission rate TR.** It combines estimations and uncertainties of within-host viral load $V_{host}$, probability of virus-positive mosquito legs $P_{leg}$, virus titer in mosquito legs $V_{leg}$, and probability of virus-positive mosquito saliva $P_{saliva}$.

All computational work presented in the manuscript was performed using R version 4.4.2 [1].

### 1 Within-host viral dynamics

To avoid converging to a local maximum, we used two sets of starting values and checked that they converged to similar values.

set 1 : $V_p$= 5.5, $T_p$= 4, $l_g$= 4.2, $l_d$= 4.5

set 2 : $V_p$= 4.5, $T_p$= 5, $l_g$= 3.2, $l_d$= 5.5

For each set, all three experimental groups were tested as the reference group. The model fit yielding the minimum -2.log-likelihood was obtained when using squirrel monkeys infected with ZIKV sylvatic strain as the reference group, and the first set of starting values. The remaining settings were $k_1$= 400, $k_2$= 150, MCMC chains = 3. The identity matrix was used as the covariance model.

The uncertainty around individual fits was obtained by sampling 500 parameter values in a multinormal distribution. As a variance-covariance matrix, we used a diagonal matrix with the estimate of the variance of the individual (cond.var.phi in saemix [2]). The 2.5% and 97.5% percentiles were then extracted from the corresponding trajectories.

The uncertainty around population-level fits, for each experimental group, was obtained with a similar procedure. Two different variance-covariance matrices were used, depending on whether the uncertainty reflected the estimation of the fixed effects, or the inter-individual variability within the experimental group (random effects). For the fixed effects, we first computed $\frac{1}{n}FIM^{-1}$, with $n$ the total number of individuals, and $FIM$ the Fisher information matrix. We then select a subset of this matrix (12x12), corresponding to parameters $V_p$, $T_p$, $l_g$, $l_d$, as well as $\beta_{species}$ and $\beta_{strain}$ for each of these four parameters. For the random effects, we use the variance-covariance matrix of the random effects (omega in saemix).

Table S 1: **Results of the non-linear mixed effect model of within-host viral dynamics, testing the effect of monkey species, viral strain, and dose delivered to the monkeys by infecting mosquitoes.** Parameter estimates (relative standard error, %) *p value* (for covariates species, strain, and dose). Squirrel monkeys infected with the sylvatic strain of ZIKV are taken as the reference.

| Parameter | Fixed Effect | Random effect SD |
|---|---|---|
| $V_p$ | 6.48 (64.8) | 0.059 (38) |
| $\beta_{species,V_p}$ | 0.13 (180.3) *0.58* | |
| $\beta_{strain,V_p}$ | -0.55 (49.1) *0.042* | |
| $\beta_{dose,V_p}$ | -0.044 (420.3) *0.81* | |
| $T_p$ | 2.63 (30.6) | 0.0046 (49) |
| $\beta_{species,T_p}$ | 0.083 (110.6) *0.37* | |
| $\beta_{strain,T_p}$ | 0.39 (23.9) *2.8e-05* | |
| $\beta_{dose,T_p}$ | 0.14 (61.5) *0.10* | |
| $l_g$ | 14.41 (89.0) | 0.11 (38) |
| $\beta_{species,l_g}$ | 0.74 (43.9) *0.023* | |
| $\beta_{strain,l_g}$ | -0.72 (50.9) *0.049* | |
| $\beta_{dose,l_g}$ | -0.40 (62.9) *0.11* | |
| $l_d$ | 7.21 (174.7) | 0.11 (72) |
| $\beta_{species,l_d}$ | 0.37 (151.4) *0.51* | |
| $\beta_{strain,l_d}$ | 0.0091 (7373.4) *0.99* | |
| $\beta_{dose,l_d}$ | -0.051 (973.1) *0.92* | |

Table S 2: **Results of the non-linear mixed effect model of within-host viral dynamics, testing the effect of monkey species and viral strain (model presented in main text).** Parameter estimates and confidence intervals (CI), derived from fixed effects and random effects separately, for each experimental group.

| Parameter | Host species | ZIKV strain | Estimate | Fixed Effect CI | Random effect CI |
|---|---|---|---|---|---|
| $V_p$ | Squirrel monkey | Sylvatic | 5.48 | [4.56 ; 6.87] | [3.43 ; 8.96] |
| $V_p$ | Cyno. macaque | Sylvatic | 6.05 | [5.10 ; 7.44] | [3.92 ; 9.64] |
| $V_p$ | Cyno. macaque | Human | 3.90 | [3.30 ; 4.70] | [2.59 ; 6.09] |
| $T_p$ | Squirrel monkey | Sylvatic | 4.15 | [3.87 ; 4.45] | [3.49 ; 4.97] |
| $T_p$ | Cyno. macaque | Sylvatic | 4.51 | [4.23 ; 4.80] | [3.68 ; 5.30] |
| $T_p$ | Cyno. macaque | Human | 5.98 | [5.65 ; 6.35] | [5.04 ; 7.09] |
| $l_g$ | Squirrel monkey | Sylvatic | 3.71 | [3.02 ; 4.60] | [1.72 ; 7.74] |
| $l_g$ | Cyno. macaque | Sylvatic | 5.05 | [4.22 ; 6.11] | [2.44 ; 10.48] |
| $l_g$ | Cyno. macaque | Human | 3.65 | [3.00 ; 4.42] | [1.84 ; 7.47] |
| $l_d$ | Squirrel monkey | Sylvatic | 6.60 | [3.20 ; 13.27] | [2.48 ; 17.01] |
| $l_d$ | Cyno. macaque | Sylvatic | 7.75 | [4.16 ; 14.16] | [2.97 ; 17.62] |
| $l_d$ | Cyno. macaque | Human | 13.41 | [7.18 ; 24.46] | [5.20 ; 32.51] |

### 2 Vector infection

#### 2.1 Disseminated infection

All models included the three variables of interest, i.e host viremia, day post infection (dpi), and host species. The goal of model selection was indeed not to decide on the inclusion of covariates but rather on the most appropriate shape of their effect. Except host species, which always had a categorical effect, the possible functional forms tested were: linear, non-parametric (tested for dpi only as it was discrete, with each level having its own fitted effect), smooth non-linear, and shape-constrained smooth non-linear. We tried to balance three criteria, namely model fit, parsimony, and biological plausibility. For model fit, we compared models using Akaike Information Criterion (AIC) but also deviance explained and pseudo-$R^2$.

For models composed only of linear, categorial, and non-parametric effects, we used generalized linear models (GLMs) fitted using the package glmmTMB [3] which conveniently allows for a thorough inspection of model residuals using the DHARMa package [4]. If a model included a smooth non-linear effect, we used generalized additive models (GAMs) with the mgcv package [5]. If a model included a shape-constrained smooth non-linear effect, we turned to shape-constrained additive models. These can be done with the scam package [6], but because they use different methods to calculate penalization and effective degrees of freedom, the AIC of gam and scam models cannot be directly compared. We formulated shape-constrained additive models in gam first, for model selection, and then used the same model in scam for numerical results and plotting purposes. In the interest of parsimony, all non-linear smooth terms were constructed with 3 interior knots and all shape-constrained smooth terms were constructed with 4 interior knots; the minimum allowed in the respective packages.

##### 2.1.1 Probability of disseminated infection

All models were fitted on summarized experimental data, using the fraction of mosquitoes with virus-positive legs for a given monkey-day combination. The response variable was expressed as k infected mosquitoes out of n tested, with k and n specific to each monkey-day combination. For this step, all data up to day 8 (last day with detected disseminated infection) was used.

It was clear from the model with linear effects for both viremia and dpi (AIC = 153, 4 degrees of freedom (df)) that this functional form was not adapted for dpi, as the model residuals showed a clear misfit. We tested a non-parametric effect for dpi, which significantly improved model performances (AIC = 146, df = 9). However, the variation of the effect from day to day was too irregular to be biologically plausible and was more likely a sign of overfitting. The overall tendency suggested that a bell-shaped effect would be appropriate. Indeed a gam with a spline for dpi showed similar performances (AIC = 148) with less parameters to be fitted (df = 5). Finally, we tried optimizing the functional form for viremia, first using a spline, which improved performances (AIC = 137, df = 6) and suggested that the effect was non-decreasing. Constraining the shape improved AIC (127), but was not in the interest of parsimony (df = 7), when formulated with gam. Once formulated in scam, the number of effective degrees of freedom dropped to 5. This model also had the advantage of anchoring the effect of viremia, with a probability of disseminated infection of 0 for viremia equal to 0, which was more biologically realistic but not possible using gam. We therefore selected this model for further inspection. We note that the significance of dpi varied from one model to the next, and even among models with a non-linear, bell-shaped effect of dpi, 2 out of 4 models tested estimated this effect to be significant.

##### 2.1.2 Viral titer in mosquito legs

For this step, only positive leg titers were used for model fitting, as we aimed to characterize what influences the value of the leg titer once we know it is positive. The simplest linear model was selected, in which only viremia had a significant effect (p = 7.6e-10). For each $\log_{10}$-unit increase in viremia, the $\log_{10}$ ZIKV titer in mosquito legs ($V_{leg}$) increased by 0.71 (95% confidence interval [0.49 ; 0.94]).

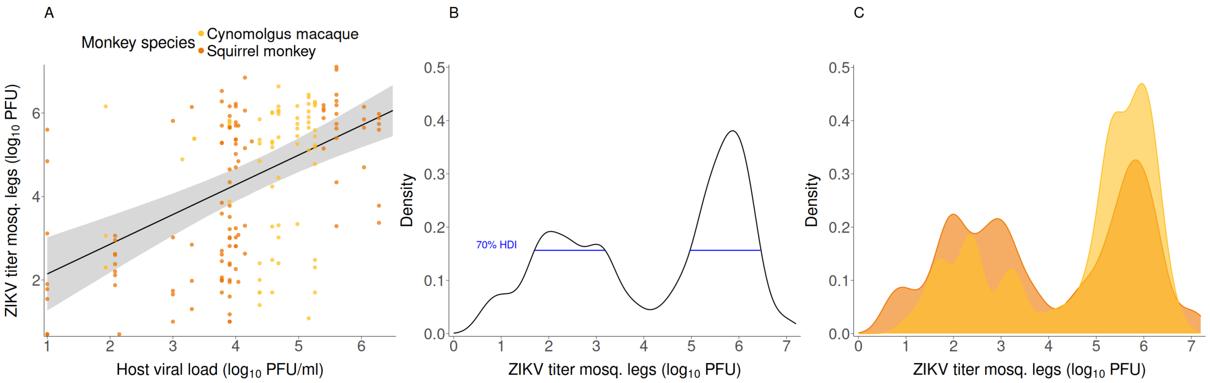


Figure S 3: A - Linear relationship between host viral load and ZIKV titer in mosquito legs. Points show raw data and the line and band show the marginal effect of viremia and its 95% confidence interval. The marginal effect of viremia represents its impact on the response variable while averaging over the effects of day of infection and monkey species, which are also included in the underlying model. Instead of fixing these variables at specific values, the marginal effect accounts for their distribution in the dataset, providing a more general view of how viremia influences the outcome across all observations. B - Empirical distribution of ZIKV titer in mosquito legs, from both monkey species. This distribution was obtained by sampling 10,000 values in the original list of measured titers, with replacement. Blue segments represent the 70% highest density interval (HDI), which is in two parts, indicating a bimodal distribution. C - Empirical distribution of ZIKV titer in mosquito legs, from each monkey species.

#### 2.2 Probability of ZIKV presence in saliva

We fitted three different functional forms (Equations S1-S3) to the data, using a maximum likelihood approach. In these equations, $p$ stands for the probability of mosquito saliva testing positive for ZIKV and $V$ is the infectious titer measured in mosquito legs, on a linear scale. Each functional form was fitted with either a binomial likelihood or a beta-binomial likelihood, the latter accounting for overdispersion in the data. We used corrected AIC (AICc) to select the functional form providing the best fit to data.

$$p(V) = \frac{1}{1 + \exp(-\beta_1(\log_{10}(V) - \log_{10}(\beta_0)))} \tag{S1}$$

$$p(V) = 1 - \exp(-(\frac{\log_{10}(V)}{\theta_0})^{\theta_1}) \tag{S2}$$

$$p(V) = \frac{\log_{10}(V)^{\gamma_1}}{\gamma_0 + \log_{10}(V)^{\gamma_1}} \tag{S3}$$

Equation S1 is the logistic function, with the curve's maximum value fixed at 1 to be on the scale of probabilities. Equation S2 was used to study vector competence for dengue when carrying Wolbachia [7]. Equation S3, also called the Hill equation, is often used to model biological interactions that demonstrate sigmoidal response, particularly to capture the biomolecular interaction exhibiting cooperativity among two binding molecules [8].

The uncertainty around the fit was obtained by sampling 500 parameter values in a multinormal distribution. We used the variance-covariance matrix from the selected model. The 2.5% and 97.5% percentiles were then extracted from the corresponding trajectories.

#### 3 Host infectiousness over time

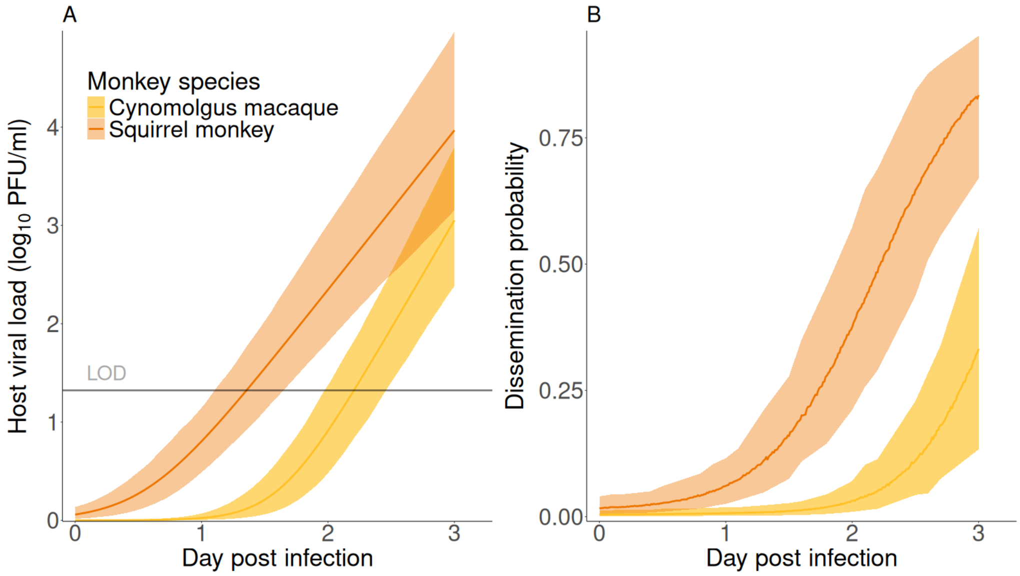


Figure S 4: **Components of monkey infectiousness in the early days of infection.** A - Monkey viral load. B - Dissemination probability. LOD = limit of detection.

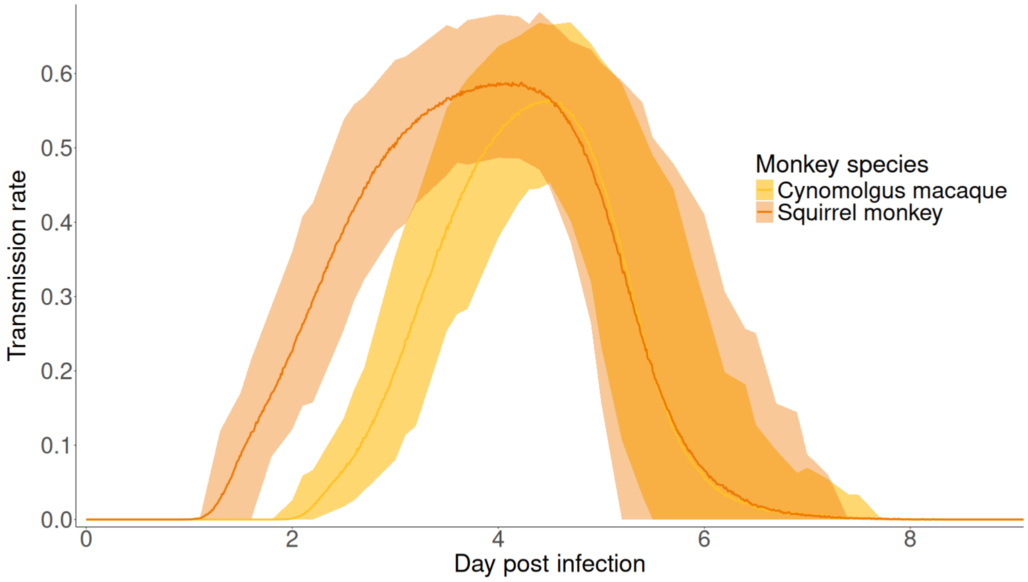


Figure S 5: **Infectiousness of squirrel monkeys and cynomolgus macaques towards *Ae. albopictus* over time, neglecting transmission from viral loads below the limit of detection.** Lines show the mean transmission rate (probability of mosquito saliva testing positive for ZIKV over the total number of mosquitoes fed) and shaded bans are the associated 95% confidence interval.

### Credits

The icon showing the mosquito with virus was downloaded for free from Flaticon and was created by Leremy : https://www.flaticon.com/authors/leremy and type “moustique” in the search bar.

The icon showing the mosquito with stripes was downloaded for free from Flaticon and was created by Surang : https://www.flaticon.com/authors/surang and type “mosquito” in the search bar. It was modified to highlight the legs or add a drop of saliva.

The icon showing the syringe was downloaded for free from Flaticon and was created by Iconfield : https://www.flaticon.com/authors/iconfield and type “syringe” in the search bar.

### References


1. R Core Team. In press. *R: A language and environment for statistical computing.* Vienna, Austria: R Foundation for Statistical Computing. See https://www.R-project.org/.

2. Comets E, Lavenu A, Lavielle M. 2017 Parameter estimation in nonlinear mixed effect models using saemix, an r implementation of the SAEM algorithm. *Journal of Statistical Software* **80**, 1–41. (doi:10.18637/jss.v080.i03)

3. Brooks ME, Kristensen K, Benthem KJ van, Magnusson A, Berg CW, Nielsen A, Skaug HJ, Mächler M, Bolker BM. 2017 glmmTMB balances speed and flexibility among packages for zero-inflated generalized linear mixed modeling. *The R Journal* **9**, 378–400.

4. Hartig F. 2022 *DHARMa: Residual diagnostics for hierarchical (multi-level / mixed) regression models*. See https://CRAN.R-project.org/package=DHARMa.

5. Wood SN. 2017 *Generalized additive models: An introduction with r, second edition.* Chapman; Hall/CRC.

6. Pya N, Wood SN. 2015 Shape constrained additive models. *Statistics and Computing* **25**, 543–559.

7. Ferguson NM *et al.* 2015 Modeling the impact on virus transmission of *Wolbachia* -mediated blocking of dengue virus infection of *Aedes aegypti*. *Science Translational Medicine* **7**, 279ra37–279ra37. (doi:10.1126/scitranslmed.3010370)

8. Somvanshi PR, Venkatesh KV. 2013 Hill equation. In *Encyclopedia of systems biology* (eds W Dubitzky, O Wolkenhauer, KH Cho, H Yokota), pp. 892–895. Springer New York.
